# High-Sensitivity Magnetic Levitation Reveals Intrinsic Protein Corona Heterogeneity on Identical Nanoparticles

**DOI:** 10.64898/2026.05.23.727410

**Authors:** Samantha Velazquez, William Thompson, Ali Akbar Ashkarran

## Abstract

The protein corona (PC), a layer of biomolecules that adsorbs onto nanoparticles (NPs) surfaces upon exposure to biological fluids, plays a key role in defining the biological identity and performance of nanomaterials. However, most current analytical approaches rely on pooled measurements of PC-coated NPs and therefore lack the resolution needed to detect subtle heterogeneity in PC composition, potentially masking important differences in NPs biological identity. Here, we used a high-sensitivity magnetic levitation (MagLev) platform capable of resolving extremely small density differences among nominally identical PC-coated NPs, enabling fractionation of particles based on subtle variations in PC composition. Compared with conventional standard MagLev systems (density resolution ∼10^-3^ g/cm^3^), the high-sensitivity MagLev improves density sensitivity by up to three orders of magnitude, allowing discrimination of density differences as small as 10^-5^ g/cm^3^. Using this approach, PC-coated NPs were separated along the MagLev column into multiple fractions corresponding to distinct density populations. Subsequent proteomic analysis across the extracted fractions identified more than 500 proteins and revealed a structured but continuous redistribution of protein composition across the column, including fraction-dependent differences in protein abundance, overlap, and biological identity. In particular, the fraction series captured hidden heterogeneity among nominally identical PC-coated NPs, with upper fractions retaining relatively stronger extracellular/plasma-associated signatures and lower fractions showing increasing representation of structural, membrane-associated, cytoskeletal, and metabolic proteins. These findings demonstrate that PC formation is intrinsically heterogeneous even on identical NPs and this heterogeneity is largely missed by conventional pooled analysis. High-sensitivity MagLev provides a simple, label-free framework for resolving PC heterogeneity and offers a new analytical approach for studying NP–biomolecule interactions, with important implications for nanomedicine design, biomarker discovery, and the clinical translation of NP-based systems.

**TOC:** 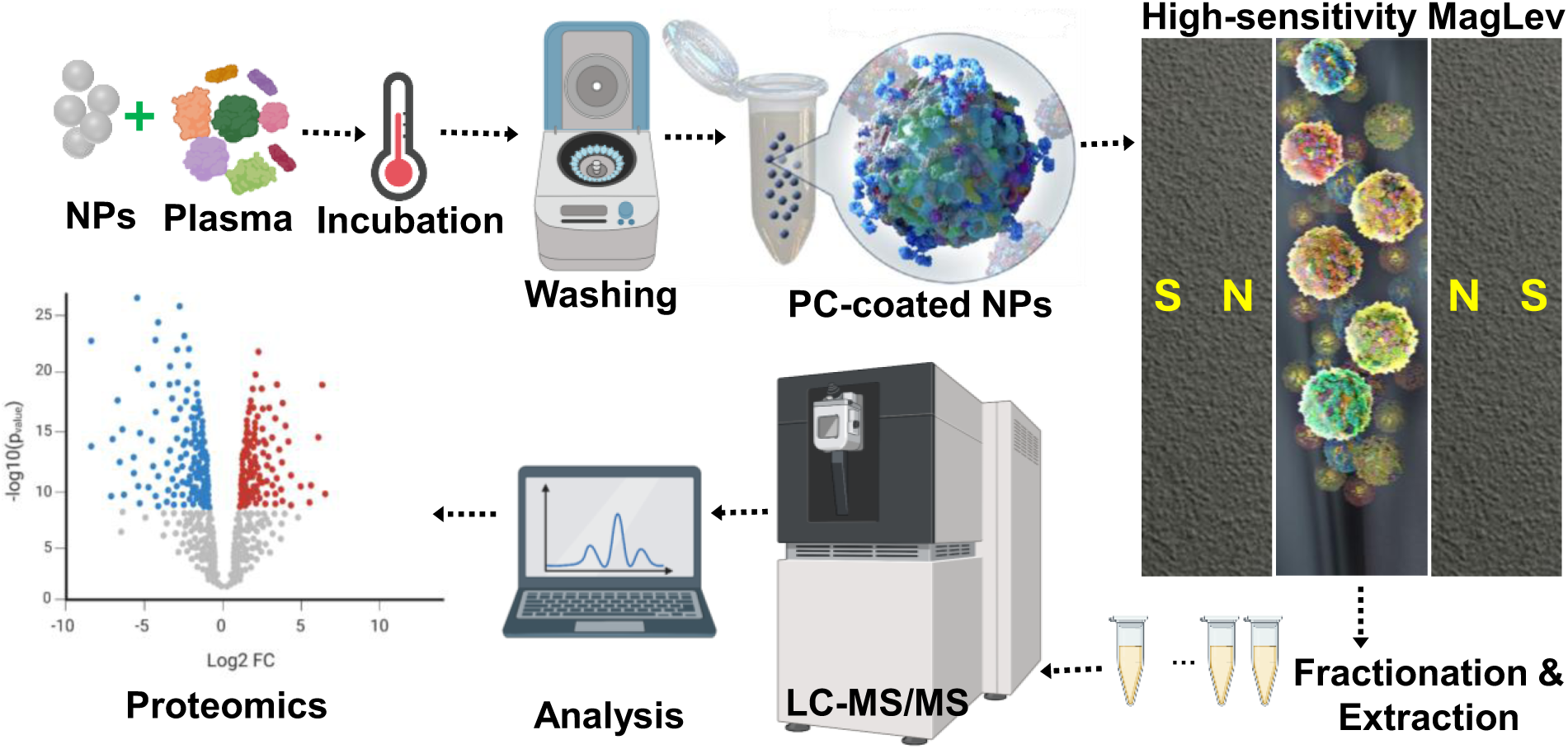

## Introduction

The PC is a complex layer of biomolecules (mainly proteins) that forms around the surface of nanoparticles (NPs) after contact with a biomolecular source such as biological fluids (e.g., blood or serum).^1–4^ The evolution of PC *in vivo* still remains highly challenging due to the complexity of human physiology and the complex nature of the biomolecules adsorbed on the surface of the NPs.^5–8^ The composition of the PC around the NPs can be highly complex and heterogeneous, meaning that the type and number of proteins that bind to the surface of the nanoparticles can vary from particle to particle, even within a single biological sample (e.g., single human plasma) and identical NPs (e.g., commercial polystyrene NPs).^9–11^ Furthermore, the recently reported personalized PC due to the variations in human proteome (e.g., in the early stages of most chronic diseases) adds more to the complex nature of the human plasma and subsequently composition and heterogeneity of PC.^12–13^

The *ex vivo* approaches for preparation of PC on the surface of model NPs do not provide homogenous PC formation as commonly perceived in the literature and the composition of the PC varies even across identical NPs with the same sizes in the same batch.^14^ The current conventional available PC characterizations or isolations techniques (e.g., liquid chromatography mass spectroscopy (LC-MS)) are not able to detect small variations in the compositions of the PC and heterogeneity of the PC structure.^15–16^ Non-specific extraction and pool analysis of the PC composition in ex vivo condition using LC-MS can mask important heterogeneities in the PC composition and may cause error and/or misunderstanding of the PC outcomes.^17–20^ The ability to distinguish extra small differences in heterogeneity and composition of the PC is crucial not only in nanomedicine research but also for clinical translation of nanomaterials.

Magnetic levitation (MagLev) has emerged as promising technique for accurate and robust density-based separations and measurements of a wide range of diamagnetic materials including various biomolecules.^21–30^ It has been used for a wide range of biological applications from cell sorting and tissue engineering to bone regeneration and self-assembly of living materials in the past years.^31–37^ Using a standard MagLev system, we were able to detect airborne viruses by concentrating the cells on the MagLev platform, extracting the viruses and their corresponding RNA, performing PCR, and identifying the viral species.^38^ There is a growing interest in using MagLev in cell biology and other novel applications as a label-free technology with a disease detection capacity. ^25, 28, 30, 39–45^ For example, in the case of sickle-cell disease, there are density differences between sickled and normal erythrocytes, which enabled a MagLev system to be used as a point-of-care diagnostic device.^46–49^ A MagLev system has been also used for differentiation between healthy and cancerous cells (i.e., circulating tumor cells) mainly due to differences in their cellular densities.^29, 32^ It is also reported that MagLev enables label-free detection of lipid accumulation in cells, offering a novel biophysical tool for disease-related cellular phenotyping.^50^ We have recently introduced MagLev as a technology for disease discrimination using pattern recognition combined with omics analysis of the levitated human plasma patterns.^51–53^ We have been able to levitate human plasma samples in MagLev system using superparamagnetic iron oxide nanoparticles (SPIONs) and uncovered that the levitated plasma carry information about the health spectrum of plasma donors.^52–53^ We have also shown that MagLev is able to discriminate PC-coated NPs from uncoated particles after exposure to human plasma.^38^ In conventional configurations (i.e., standard MagLev systems), however, its resolving power is limited and may not be sufficient to distinguish extremely small differences among nominally identical PC-coated NPs.^54^ For this reason, there remains a need for higher-sensitivity approaches capable of resolving fine heterogeneity that is otherwise hidden in bulk measurements.^55^ Therefore, a method that can separate closely related NPs subpopulations before downstream biochemical analysis would provide a more realistic picture of PC organization and would help connect physical fractionation with molecular composition.^56–58^

To address this challenge, we used a high-sensitivity MagLev system rather than a conventional MagLev configuration in order to detect subtle differences in PC composition and reveal hidden heterogeneity among nominally identical PC-coated NPs. In this study, we incubated commercially available polystyrene NPs (∼90 nm) with human plasma, followed by isolation and purification of the resulting PC-coated NPs. The PC-coated NPs were then introduced into a high-sensitivity MagLev system containing SPIONs, and their levitation profiles were monitored over a period of 2 h. At the end of levitation, fractions were collected and analyzed by sodium dodecyl sulfate-polyacrylamide gel electrophoresis (SDS-PAGE) and liquid chromatography-mass spectrometry (LC-MS/MS). The goal of this work was to develop an approach for detecting and resolving hidden heterogeneity in NPs’-PC and to determine whether nominally identical PC-coated NPs contain compositionally distinct subpopulations that are missed by conventional pooled analysis. By combining high-sensitivity MagLev with fraction-resolved proteomics, this study provides a new way to probe PC organization at the nano-biointerface and offers insight into how subtle compositional differences may influence the biological identity of NPs.

### Experimental details

#### Materials

SPIONs (30 mg/mL) functionalized with ferumoxytol were purchased from Feraheme (www.feraheme.com) and diluted with phosphate buffered saline (PBS, 1X) solution to desired concentrations. Fluorescent polyethylene microparticles with known densities were obtained from Cospheric (www.cospheric.com). Pooled healthy human plasma samples ordered from Innovative Research and were diluted to a final concentration of 55% using PBS, 1X. Plain polystyrene NPs (∼ 90 nm) were provided by Polysciences. (www.polysciences.com). Ammonium bicarbonate, urea, dithiothreitol, iodoacetamide, and acetonitrile were purchased from Sigma-Aldrich. Trypsin was obtained from Promega, and formic acid was purchased from Thermo Scientific.

#### MagLev platform

A high-sensitivity MagLev system was constructed using two opposing rare-Earth 6 in × 2 in × 1 in N42 neodymium magnets (model NB085-5, Magnet4less). The magnets were arranged with like poles facing each other in a 90° rotated configuration (**Figure 1**), producing a strong magnetic-field gradient along the horizontal axis and a weaker magnetic-field gradient along the vertical axis. In this geometry, the strong horizontal gradient helps focus the sample toward the center, whereas the weaker vertical gradient, acting together with gravity, promotes separation along the column and increases sensitivity.^55^ The levitation behavior and overall resolution of the system depend strongly on three main parameters: the distance between the magnets, the magnetic field strength, and the concentration of the paramagnetic medium. The two magnets were mounted in custom 3D-printed holders fixed onto an aluminum breadboard using posts, bearings, support beams, and a dual lead ball screw assembly, which allowed the magnet spacing to be adjusted easily using a hand crank (**Figure 1**). The sample holder was also 3D-printed and positioned between the magnets using a dedicated support structure. We also used a custom-built test tube to minimize the amount of paramagnetic liquid required for levitation experiments. In contrast to a conventional MagLev setup, the platform was intentionally operated in a rotated geometry in which the stronger magnetic-field gradient was oriented predominantly along the horizontal axis, while the weaker gradient was aligned with gravity along the vertical axis. In fact, the strong horizontal gradient provided lateral focusing and stability, whereas the weak vertical gradient increased the spatial separation of levitating populations with very small effective density differences. As a result, this geometry improved the sensitivity of the platform for resolving subtle heterogeneity among nominally identical PC-coated NPs, as previously reported.^55^ Optical images and levitation profiles were recorded using a Nikon D750 digital camera equipped with a 105 mm Nikkor Micro lens, together with a ruler carrying millimeter-scale markings for position measurements.

**Figure 1.**
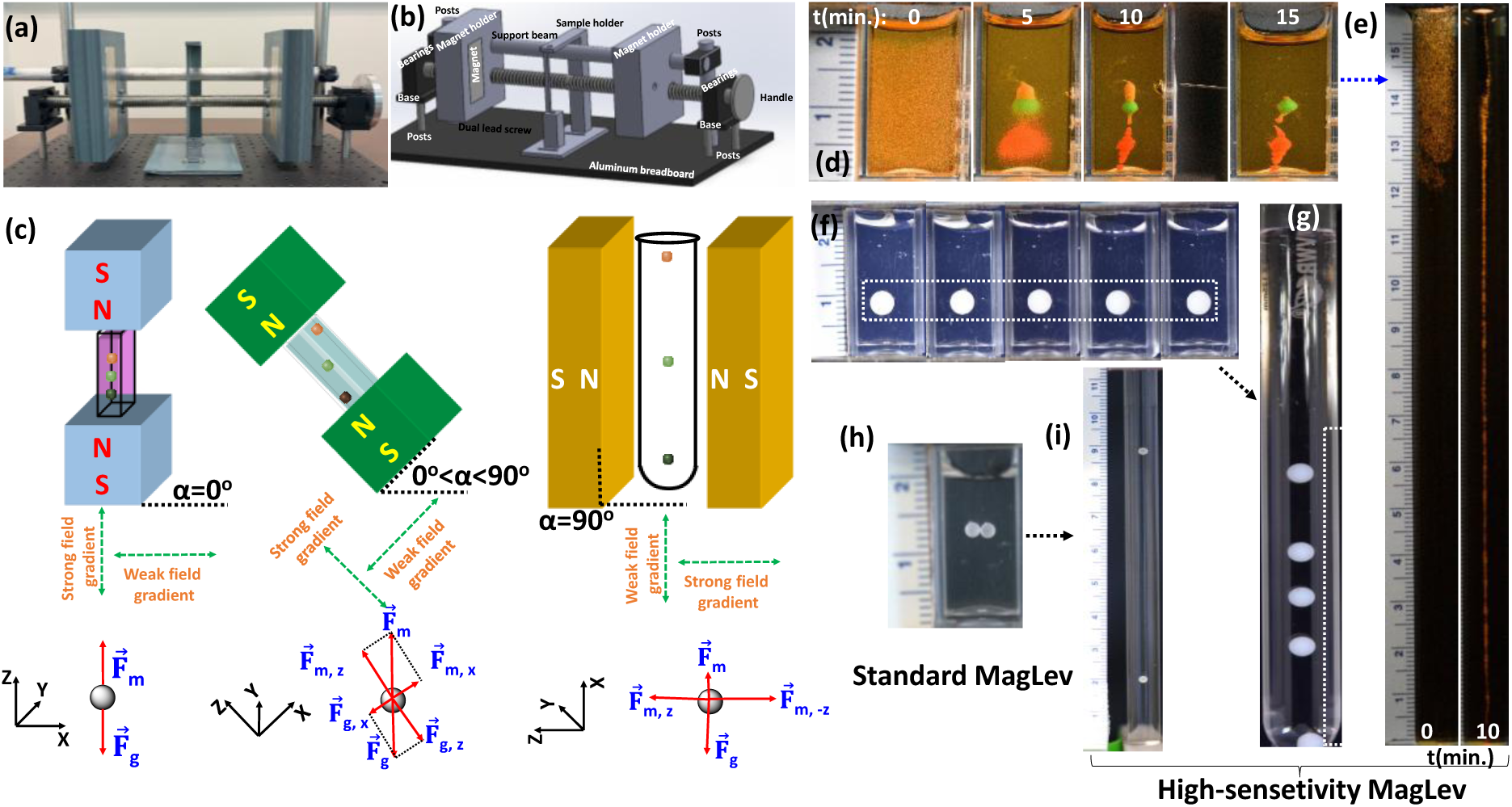
(a) Photograph and (b) schematic of the high-sensitivity MagLev configuration used in this study, (c) schematic showing how the resolution increases after 90° rotation compared with the standard MagLev configuration, (d) levitation profiles of mixed fluorescent microspheres with different densities (∼40–50 µm in size) in SPIONs (0.125mg/ml), (e) levitation profiles of 1.011 g/cm³ particles in the high-sensitivity MagLev after extraction from the standard MagLev in SPIONs (0.125mg/ml), demonstrating how a few millimeters of levitation distance can be translated into more than 10 cm due to the increased sensitivity, (f) levitation of five commercial nylon beads (∼ 4mm in size and density of 1.14g/cm^3^) from the same batch, and (g) the corresponding levitation of the same five nylon beads in the high-sensitivity MagLev, (h) levitation of two identical commercial polystyrene beads (∼ 3mm in size and density of 1.05g/cm^3^), and (i) the corresponding levitation of the same two polystyrene beads in the high-sensitivity MagLev system.

#### PC formation on the surface of the NPs

For PC formation, NPs were incubated with diluted (55%) plasma samples from healthy individuals so that the NPs’ and human plasma final concentrations were 0.2 mg/ml and 55% in a 1.5 ml batch, respectively. The solution was then mixed, vortexed well, and incubated for 1h at 37 °C at a constant 1000 rpm agitation. To remove unbound and plasma proteins only loosely attached to the surface of NPs, protein-NPs complexes were then centrifuged at 17,000g for 20 minutes, the collected NPs’ pellets were washed twice more with cold PBS under the same conditions, and the final pellet was re-dispersed in 50 uL of PBS for injection into the high-sensitivity MagLev system.

#### Magnetic levitation fractionation and sample collection

After injection of the PC-coated NPs into the high-sensitivity MagLev system and the levitation experiments were initiated, the PC-coated NPs were allowed to reach their equilibrium levitation distribution within the MagLev column. Based on time-course observations, the levitation profiles were completely stabilized after about 2 h, with no significant change in levitation height thereafter. Once the levitation pattern reached equilibrium, the levitated cloud of PC-coated NPs was fractionated along the vertical axis of the MagLev column. Fractions were collected sequentially at 1 cm vertical intervals along the levitation column, beginning from the top of the column and proceeding downward. Visual inspection of the levitation profiles indicated that the majority of the particle population was distributed between 14 cm and 3 cm along the column. Regions above 14 cm and below 3 cm contained negligible particle populations and were therefore excluded from further analysis. Consequently, the MagLev column was divided into discrete fractions corresponding to levitation heights between 14 cm and 3 cm, generating a series of height-resolved samples representing particles with slightly different effective densities. Each fraction was carefully extracted layer-by-layer using a micropipette (**Figure S1**) to minimize disturbance of adjacent regions. The collected fractions were transferred into individual low bindings Eppendorf tubes and centrifuged at 17000g for 20 min to collect the PC-coated NPs from the SPIONs solution for further analysis.

#### LC-MS/MS sample preparation

The collected NPs’ PC pellets in the previous step (i.e., after fractionation by high-sensitivity MagLev system) were subjected to a standardized proteomic digestion protocol optimized for mass spectrometry based on our previous findings.^59^ Each PC-coated NPs’ pellet was resuspended in 25 µL of buffer containing 50 mM ammonium bicarbonate and 1.6 M urea to facilitate protein solubilization and denaturation. Reduction was achieved by adding 2.5 µL of dithiothreitol (DTT, Pierce BondBreaker, neutral pH) to a final concentration of 10 mM. Samples were incubated at 37 °C for 60 minutes using a Thermomixer. Following incubation, samples were brought to room temperature, vortexed briefly, and centrifuged to collect the contents. Alkylation was performed by adding 2.7 µL of freshly prepared iodoacetamide (IAA) to a final concentration of 50 mM followed by incubation in the dark at room temperature for 1 hour. To quench excess IAA, 3 µL of DTT (10 mM final) was added, followed by a 15-minute incubation at room temperature. Subsequently, sequencing-grade trypsin (Promega), diluted in protein digestion buffer, was added at a 1:50 trypsin-to-protein ratio. Enzymatic digestion proceeded overnight (∼16 h) at 37 °C in a Thermomixer. The next day, the digested samples were cooled to room temperature and centrifuged at 17,000 × g for 20 minutes to pellet residual NPs. Supernatants containing the tryptic peptides were carefully transferred into fresh low-binding Eppendorf tubes. Formic acid (FA) was added to a final concentration of 5% (v/v) to acidify the samples and adjust pH to between 2 and 3, ensuring peptide stability and compatibility with desalting. Peptide desalting and cleanup were performed using C18 StageTips. Each tip was first activated with 80% acetonitrile (ACN) and 5% FA, equilibrated with 5% FA, and then loaded with the peptide-containing samples. Bound peptides were washed with 5% FA and eluted in two stages: first with 30% ACN + 5% FA, and then with 80% ACN + 5% FA. Both eluates for each sample were pooled, gently mixed, and transferred into a new low-binding tube. Samples were dried using a vacuum centrifuge (SpeedVac) and stored at –20 °C prior to shipment. Dried peptides were shipped overnight in dry ice to the core proteomics facility using FEDEX with next-morning delivery. Sample integrity upon arrival was confirmed by the receiving facility.

#### Physicochemical characterizations

DLS and zeta potential analyses were performed to measure the size distribution and surface charge of the NPs before and after PC formation as well as after fractionation by high-sensitivity MagLev system using a Zetasizer Lab series DLS instrument (Malvern company). A Helium Neon laser with a wavelength of 632 nm was used for size distribution measurement at room temperature. TEM was carried out using a JEM-120i (JEOL Ltd.) operated at 120kV. 20 μl of the bare NPs were deposited onto a copper grid and used for imaging. For PC-coated NPs, 20 μl of sample was negatively stained using 20 μl uranyl acetate 1%, washed with DI water, deposited onto a copper grid, and used for imaging. PC composition was also determined using SDS-PAGE and LC-MS/MS analyses.

#### LC-MS/MS analysis and proteomics data processing

Each dried vial was reconstituted with 22 µl 2% ACN 0.1% FA. 5 µL of each digest was injected and run 3 times by nanoLC-MS/MS using a 2h gradient on a Waters CSH 0.075mm x 250mm C18 column coupled to a Thermo Orbitrap Astral mass spectrometer operated with HCD fragmentation in data-dependent acquisition (DDA) mode. Protein identification and primary quantification were performed in Proteome Discoverer 3.2 using Sequest HT against the UniProt human canonical database, assuming tryptic digestion. Search parameters included a 10 ppm precursor mass tolerance and 0.6 Da fragment mass tolerance; carbamidomethylation of cysteine was set as a fixed modification, and deamidation of asparagine/glutamine and oxidation of methionine were set as variable modifications. Peptide-spectrum matches were validated using Percolator, and identifications were filtered at 1% false discovery rate (FDR). Protein quantification was based on precursor ion intensity, normalized by total peptide amount, and only proteins identified with at least 2 peptides and 3 peptide-spectrum matches (PSMs) were reported. All downstream proteomics and statistical analysis were performed from the final processed 3-replicate protein report provided by the core facility (**Table S2**). For each reported comparison, the replicate abundance-ratio values were averaged and used for downstream visualization. Fraction-level protein abundance profiles were reconstructed from the fraction-to-control ratio columns, and positive ratio values were log2-transformed prior to analysis. Clustered heatmaps were generated from row-standardized protein abundance matrices, shared-protein heatmaps were generated from the pairwise overlap counts reported in the processed summary output, Principal Component Analysis (PCA) was performed on reconstructed fraction-level protein profiles after row-wise minimum-value imputation and feature standardization, and the top-changing proteins were ranked using a composite trajectory score based on endpoint difference, abundance range, and cumulative stepwise change across fractions. Replicate reproducibility was assessed from the 3-replicate outputs using protein-level coefficients of variation, and adjacent-fraction volcano plots were generated from replicate ratio values after log2 transformation and one-sample testing against zero.

## Results and discussion

An overview of the experimental workflow is illustrated in **Figure 2**. **Figure 3** shows the levitation profiles of PC-coated NPs and uncoated NPs inside the high-sensitivity MagLev system, as well as outside the MagLev system as a control at an optimized 0.125 mg/ml concentration of SPIONs. It is noteworthy that, before selecting the working SPIONs concentration, we performed a series of levitation optimization experiments over a broad concentration range (0.03–0.25 mg/mL) using both PC-coated and uncoated NPs. Based on the stability of the levitation profiles, reproducibility of the profiles, and separation sensitivity, 0.125 mg/mL was identified as the optimum SPIONs concentration for the high-sensitivity MagLev system. Therefore, all levitation profiles and fractionation results presented here were obtained using 0.125 mg/mL SPIONs, while the remaining optimization data are not shown. The time-dependent levitation profiles were recorded every 10 min for 2 h for PC-coated and uncoated NP samples inside and outside the high-sensitivity MagLev system (**Figure 3**). In general, the samples formed elongated levitation bands that gradually stabilized over time; however, clear differences were observed in both the evolution and the final shape of the levitating populations. For the PC-coated NPs inside the MagLev system (**Figure 3a**), the levitating population showed a broad and evolving profile that gradually redistributed over the course of the experiment. The band extended over a relatively large vertical region and exhibited visible changes in shape and distribution over the time, indicating the presence of heterogeneous PC-coated subpopulations with different effective levitation behaviors. This is while the same PC-coated NPs within the standard MagLev system showed a quite narrow distribution of only a few millimeters (**Figure S1**). This observation supports the main hypothesis of the study that apparently identical PC-coated NPs do not behave as a single uniform population, but instead contain compositionally distinct subgroups that can be spatially resolved by the high-sensitivity MagLev platform. In contrast, the uncoated NPs inside the MagLev system (**Figure 3b**) displayed a different levitation pattern. Although the sample still showed time-dependent redistribution, the overall levitation profile and distribution appeared different from the PC-coated sample. This difference suggests that PC formation contributes significantly to the complexity of the levitation pattern, most likely by introducing particle-to-particle differences in effective density, surface composition, and associated biomolecular structure.

**Figure 2:**
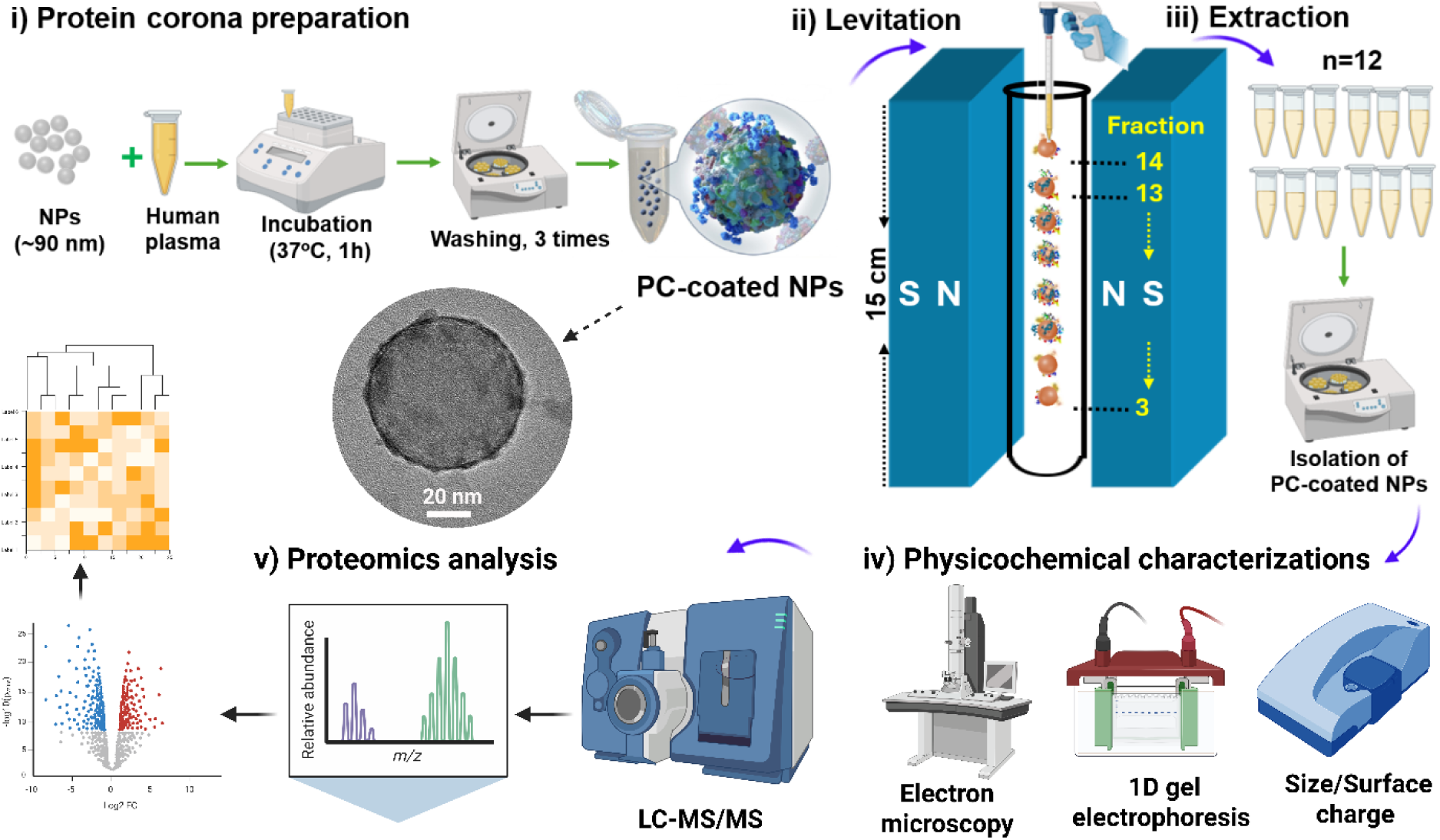
Schematic overview of the experimental workflow. Polystyrene NPs were incubated with human plasma to form PC-coated NPs, followed by injection into the high-sensitivity MagLev system and extraction of fractions from top to bottom of the MagLev column. The resulting PC-coated NPs were isolated again and subjected to physicochemical characterization, LC–MS/MS analysis, and downstream proteomics data processing.

**Figure 3:**
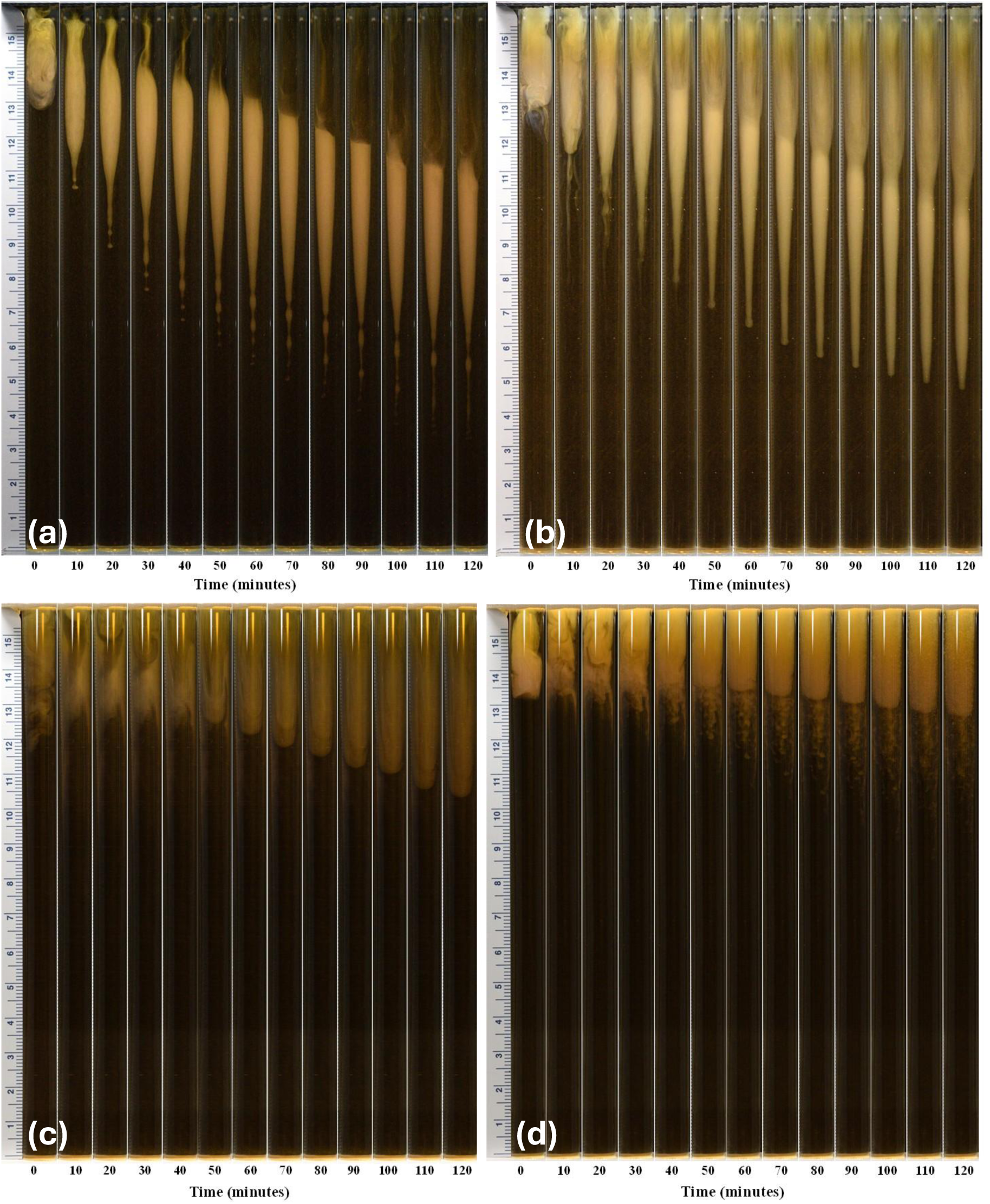
Time-dependent levitation profiles of PC-coated and uncoated NPs inside and outside the high-sensitivity MagLev system. (a) PC-coated NPs inside the high-sensitivity MagLev system, (b) uncoated NPs inside the high-sensitivity MagLev system, (c) PC-coated NPs outside the high-sensitivity MagLev system, and (d) uncoated NPs outside the high-sensitivity MagLev system.

To ensure that these behaviors occurred only inside the MagLev system and were due to the particular configuration of the MagLev, we performed the same experiments outside the MagLev system as a control. As is clear from **Figures 3c** and **3d**, the PC-coated NPs and uncoated NPs outside the MagLev system behaved totally differently and, in fact, did not show any levitation or distribution over time. Therefore, we may conclude that the levitation profiles and the corresponding variations are a unique response of PC-coated NPs with different compositions within the MagLev system.

To characterize the distribution of PC-coated NPs populations across the MagLev column, DLS analysis was performed on the collected fractions. In this study, commercially available plain polystyrene NPs with an average diameter of approximately 90 nm were used as model nanomaterials for PC formation because of their well-defined physicochemical properties, good stability, and widespread use in nanomedicine research. Our prior characterization (in addition to the analysis from the vendor), confirmed that the bare PSNPs were monodisperse, with an average hydrodynamic diameter of approximately 92 nm and a surface zeta potential of about −35 mV. After incubation with human plasma, formation of the protein corona led to an increase in hydrodynamic diameter to approximately 130 nm and a shift in surface charge to about −12 mV, confirming successful PC formation (**Figure 4** and **Table 1**). DLS analysis of the MagLev-separated fractions was then carried out to find how PC-coated NPs size distributions and particle populations varied across the levitation profile in the high-sensitivity MagLev system.

**Figure 4:**
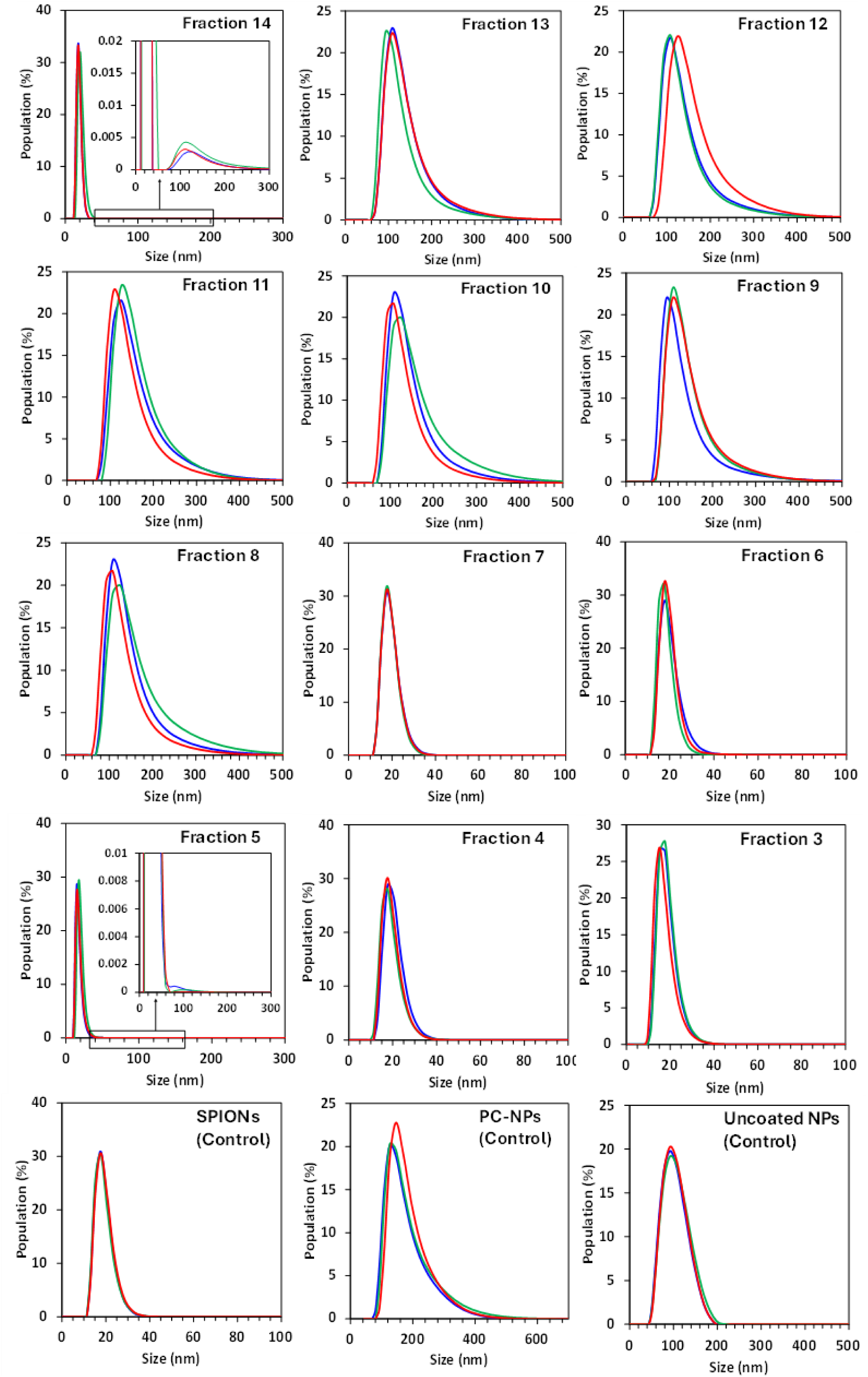
Size distributions of PC-coated NPs across high-sensitivity MagLev fractions (14–3) collected at 1 cm levitation height intervals after equilibrium (∼2 h). Each panel shows population (%) versus hydrodynamic diameter (nm) for three independent measurements (red, blue, green). Control measurements of SPIONs, PC–coated NPs, and uncoated NPs are shown as control.

**Table 1:** Average size, polydispersity index (PDI), zeta potential, and the corresponding standard deviation values of Hydrodynamic size, polydispersity index (PDI), and zeta potential of control NPs and MagLev fractions (14–3) determined by DLS measurements (three repeats).

| Sample | Size (nm) | SD (nm) | PDI | SD | Zeta potential (mV) | SD (mV) |
| --- | --- | --- | --- | --- | --- | --- |
| Uncoated NPs | 92.9 | 0 | 0.055 | 0.005 | -35.1 | 0.7 |
| PC-coated NPs | 132.4 | 9.7 | 0.354 | 0.015 | -12.3 | 0.6 |
| SPIONs | 17.7 | 0 | 0.287 | 0.023 | -16.1 | 0.9 |
| Fraction 14 | 18.6 | 1.4 | 0.310 | 0.017 | -9.5 | 0.6 |
| Fraction 13 | 103.0 | 7.1 | 0.287 | 0.017 | -12.5 | 0.7 |
| Fraction 12 | 113.9 | 8.2 | 0.244 | 0.009 | -12.4 | 0.5 |
| Fraction 11 | 119.7 | 8.3 | 0.240 | 0.007 | -13.8 | 0.4 |
| Fraction 10 | 113.9 | 8.2 | 0.240 | 0.012 | -12.1 | 0.5 |
| Fraction 9 | 103.0 | 7.1 | 0.213 | 0.012 | -11.4 | 0.9 |
| Fraction 8 | 119.7 | 8.3 | 0.233 | 0.024 | -8.6 | 0.9 |
| Fraction 7 | 17.7 | 0 | 0.592 | 0.105 | -7.4 | 0.4 |
| Fraction 6 | 17.7 | 0 | 0.430 | 0.035 | -3.3 | 0.3 |
| Fraction 5 | 16.0 | 1.2 | 0.527 | 0.047 | -3.1 | 0.9 |
| Fraction 4 | 17.7 | 0 | 0.473 | 0.014 | -3.9 | 0.6 |
| Fraction 3 | 16.4 | 1.1 | 0.359 | 0.062 | -3.1 | 0.1 |

Across the MagLev fractions, the DLS data show a clear height-dependent change in the relative populations of PC-coated NPs recovered from the column. In the very top fraction (i.e., fraction 14), the distributions are dominated by a narrow peak centered around 20 nm, consistent with the expected hydrodynamic size range of bare SPIONs NPs. As is clear from the levitation images, in the very top fraction the contribution from PC-coated NPs is relatively minor in comparison with the dominant SPIONs matrix. Moving downward through the intermediate fractions (approximately fractions 13–8), a broader peak above 100 nm appears and remains the major feature, indicating that these fractions are enriched in PC-coated NPs with hydrodynamic diameters consistent with PC formation on the NPs surface. This is in good agreement with the expected average size of the NPs after PC formation, given that the nominal NPs diameter is ∼92 nm and adsorption of plasma proteins increase the apparent hydrodynamic diameter to values above 100 nm. However, by moving further down the column, another shift in the particle-size distributions was observed. In the lower fractions (particularly fractions 7–3), the relative contribution of the small-size population below ∼20 nm increases substantially, while the abundance of the broad >100 nm population decreases. In other words, in the latest fractions the small-particle peak (i.e. <20 nm) becomes dominant again, indicating that these lower fractions contain fewer PC-coated NPs and the populations of SPIONs become dominant. Therefore, the DLS results support a transition from fractions enriched in PC-coated NPs populations at higher levitation heights to fractions increasingly dominated by smaller components (i.e., SPIONs matrix) at lower levitation heights.

An important point is that although some fractions may show average particle sizes near or below ∼20 nm, this does not mean that the PC-coated NP population is absent throughout the column. In fact, these low average values are strongly influenced by the dominant sharp peak of the SPION population, which acts as the main matrix. However, we still observe traces of PC-coated NPs, although the intensity of their detected signals is significantly lower compared with SPIONs, as depicted in the inset of fraction 14 and 4, as representatives (**Figure 4**). In the main body of the levitated cloud, the dominant population remains the PC-coated NPs, characterized by hydrodynamic diameters above 100 nm, which is consistent with PC formation on NPs with an original size of approximately 92 nm. Only in the late fractions, where the population of PC-coated NPs decreases, the detected DLS signal of ∼20 nm particles become dominant again, as shown in the inset of fraction 5 as a representative example. The control measurements further support this interpretation. The SPION control exhibits a narrow distribution centered below 20 nm, confirming the expected size range of the small magnetic particles. In contrast, the PC-NPs control shows a broad distribution above 100 nm, consistent with successful PC formation and an increased hydrodynamic diameter. The uncoated NPs control also shows a broader population in the ∼100 nm range but remains distinct from the SPIONs control. These results indicate that the MagLev fractions capture a gradual change in the relative abundance of two main populations: a larger PC-coated NP population dominating much of the main levitated cloud, and a smaller-particle population that becomes increasingly prominent in the very top and lower fractions due to the presence of matrix solution (i.e., SPIONs).

The zeta potential measurements further confirm the formation and evolution of the PC across the MagLev fractions. The bare polystyrene NPs exhibited a strongly negative surface charge (∼ −35 mV), reflecting the intrinsic surface chemistry of the polystyrene NPs. After incubation with plasma, the zeta potential shifted toward less negative values (∼ −12 mV), consistent with adsorption of plasma proteins and the formation of a PC layer that partially screens the native surface charge of the NPs. This shift is also observed in the intermediate MagLev fractions (fractions 13–8), where zeta potentials remain within a similar range (approximately −6 to −14 mV), indicating that these fractions are enriched in PC–coated NPs. In contrast, the early (fraction 14) and late fractions (fractions 7–3) exhibit less negative zeta potentials (approximately −3 to −9 mV), which correlates with the increased presence of smaller SPIONs-rich populations detected by DLS. The SPIONs control itself shows a zeta potential of ∼ −16 mV, confirming that differences in particle composition and surface chemistry across fractions influence the measured electrostatic properties. Overall, the gradual shift in zeta potential across the fractions is in agreement with the redistribution of NPs populations and the varying contributions of PC–coated NPs and smaller paramagnetic particles throughout the MagLev column.

Representative TEM images of the bare NPs and PC-coated NPs were collected to confirm particle morphology and size (**Figure 5**). As expected, the commercial uncoated polystyrene NPs appear as uniform, spherical particles that are highly monodisperse, with sizes consistent with the nominal diameter (∼90 nm), in agreement with the manufacturer’s MSDS data information sheets, as well as our previous reports and prior studies. These images are provided as representative examples of the morphology of the NPs and PC-coated NPs used in this work. No significant variations in particle size or visible differences in PC coverage were observed across samples by TEM. This is expected because TEM probes individual NPs at the nanoscale, whereas the MagLev fractionation separates NP populations on a much larger scale based on bulk density differences arising from variations in PC composition. Therefore, subtle differences in PC composition that drive separation in the MagLev system are difficult to directly visualize using TEM imaging alone. These results are in agreement with the DLS findings, which showed a larger average hydrodynamic diameter for the PC-coated NPs compared with the bare NPs.

**Figure 5:**
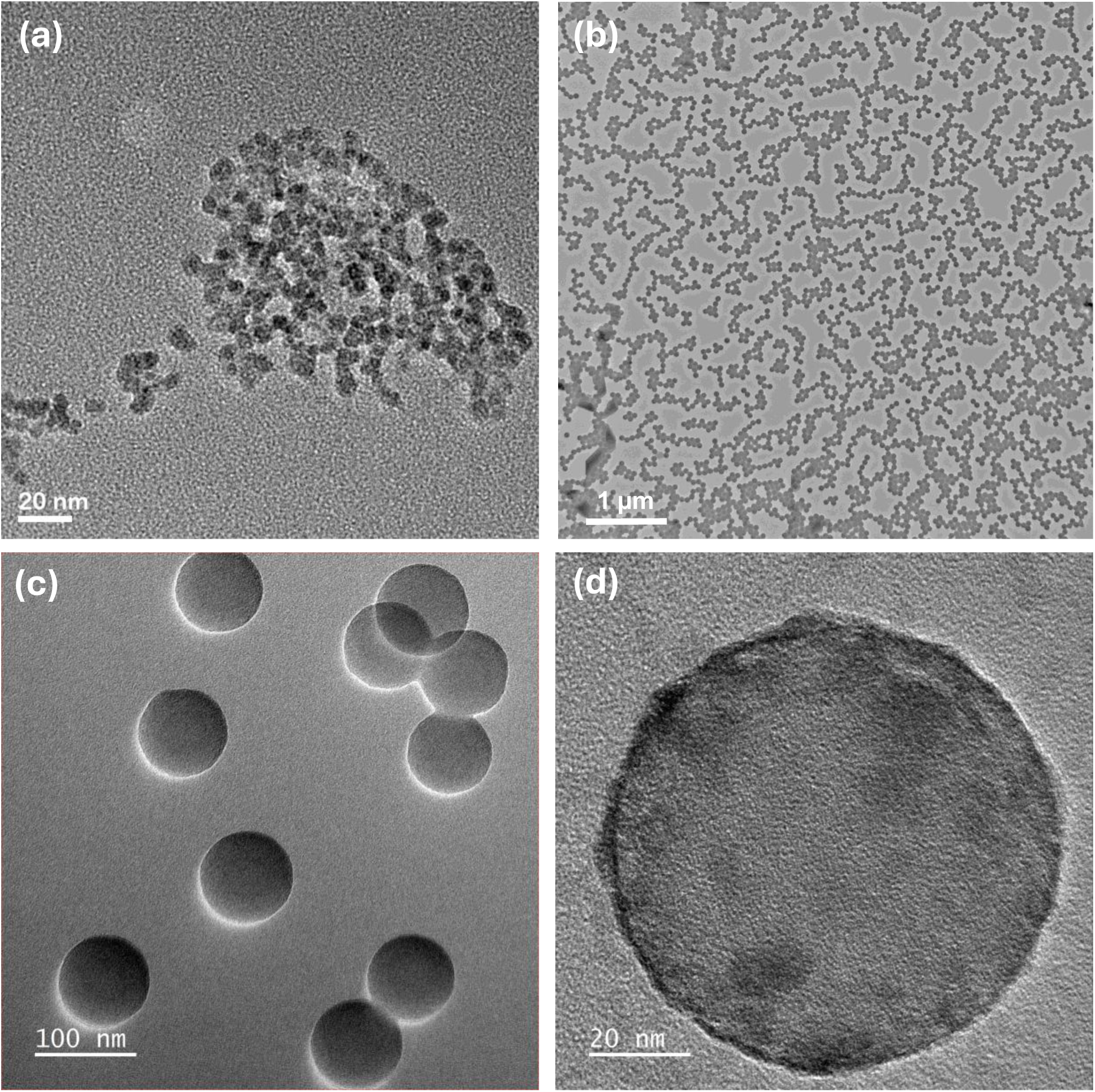
Representative TEM images of (a) bare SPIONs, (b) and (c) bare polystyrene NPs at low and high magnifications to show the monodispersity and ideal spherical shape of the NPs, and (d) formation of PC on the surface of the NPs after exposure to human plasma.

As a next step, we analyzed the MagLev-separated fractions using one-dimensional (1D) SDS-PAGE to obtain an initial view of the PC profiles across the levitation column (**Figure 6**). The plasma lane shows the expected complex mixture of abundant plasma proteins, whereas the PC-coated NPs (PCNP) lane exhibits a simplified banding pattern, indicating selective enrichment of a subset of plasma proteins on the NPs surface after PC formation. The blank lane shows no detectable bands, confirming that the observed signals arise from the nanoparticle-containing samples rather than from buffer or preparation artifacts.

**Figure 6:**
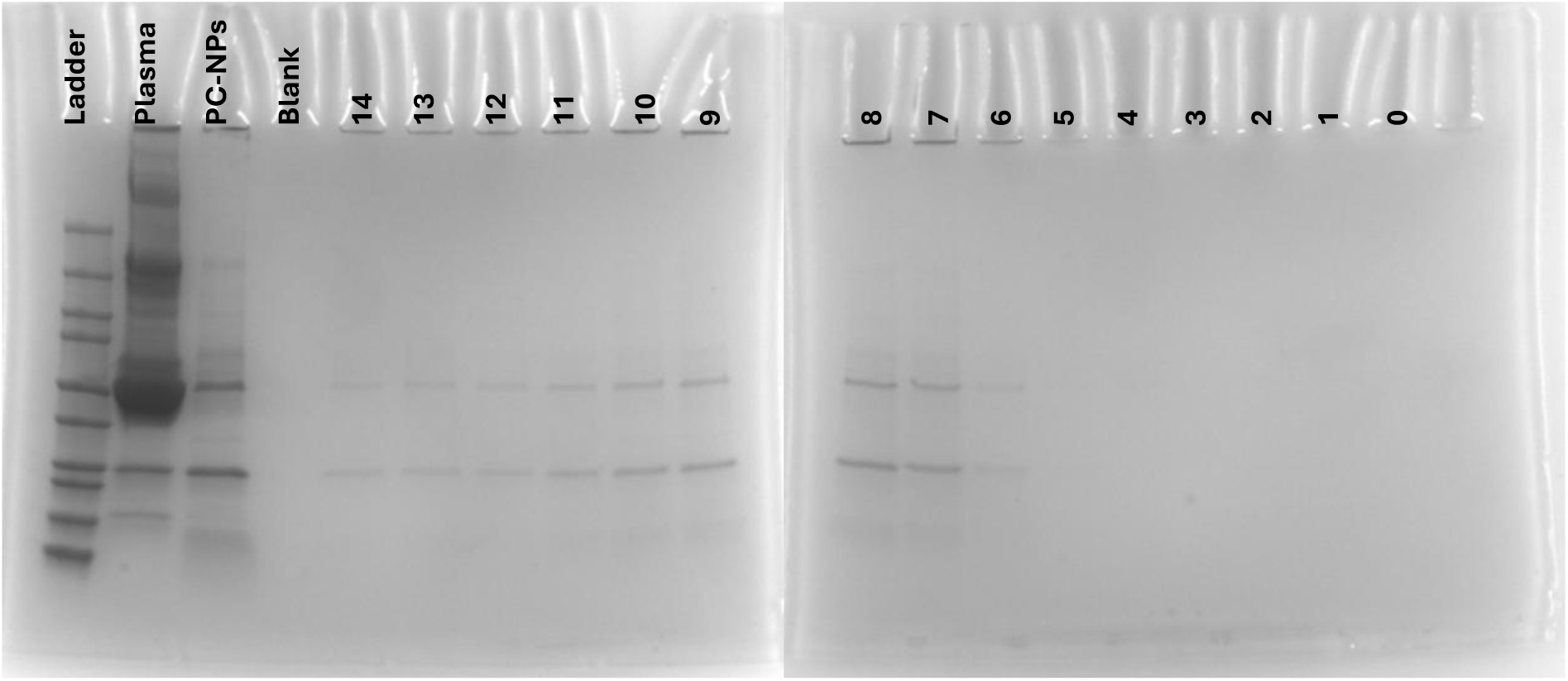
One-dimensional (1D) SDS–PAGE analysis of plasma proteins, PC-coated NPs, and MagLev-separated fractions. The gel shows the protein ladder, plasma control, PC-coated NPs as control, and fractions collected along the MagLev column from the very top (15) to the lowest part (0) of the MagLev column.

Importantly, the eluted MagLev fraction shows clear fraction-dependent differences in band intensity and distribution. The upper fractions (e.g., 15 and 14) display detectable protein bands, with the signal becoming more pronounced toward the middle of this region, while fractions 9–7 still show visible bands but with changing intensity. In contrast, the lower fractions (approximately 6–0) exhibit very weak or nearly undetectable bands. Overall, these results indicate that the PC-coated NPs are not uniformly distributed across the MagLev column, but instead are concentrated within a limited set of fractions. This observation is consistent with the optical appearance of the fractions (see for example levitation images in **Figure 3**), where the PC-coated NPs-enriched fractions appear visually distinct from the others. Overall, the SDS-PAGE results provide qualitative evidence that different MagLev fractions contain PC-coated NPs with different protein coverages and support the presence of heterogeneity in the identical NPs’ PC composition. In addition, the similarity of the dominant bands across the positive fractions suggests that 1D SDS-PAGE does not provide sufficient resolving power to fully distinguish subtle differences in PC composition between nearby fractions. Therefore, while SDS-PAGE is useful as an initial qualitative screening tool to identify protein-containing fractions and confirm non-uniform distribution across the MagLev column, higher-resolution analysis such as LC–MS/MS is required to determine the detailed compositional differences between fractions.

Next, we performed LC–MS analysis on the extracted fractions to determine the PC composition of the different PC-coated NP populations distributed across the high-sensitivity MagLev column. LC–MS analysis of the high-sensitivity MagLev fractions (**Figure 7**) showed that proteomic composition changes progressively across the column, indicating that the separation reflects a structured continuum of related protein populations. Interestingly, the total number of quantified proteins varied across the fraction series, with 497 proteins in unfractionated control, followed by 273, 277, 288, 355, 352, 392, 392, 370, 315, 259, 232, and 217 proteins in fractions 14 to 3, respectively (**Figure 7c**). This finding indicates that protein coverage does not remain constant over extended regions of the column, but instead changes continuously as the sample moves downward. In particular, proteome coverage increases from the upper fractions toward the middle region, reaches its highest values in fractions 9–8, and then gradually declines toward the lowest fractions. This trend suggests that the high-sensitivity MagLev platform resolves fine fraction-to-fraction heterogeneity that would be obscured in a lower-resolution or bulk analysis. Although sample is an ensemble of identical PC-coated NPs and would conventionally be treated as a single PC population, the high-sensitivity MagLev analysis revealed substantial heterogeneity in protein composition (e.g., in terms of number of proteins) across fractions.

**Figure 7:**
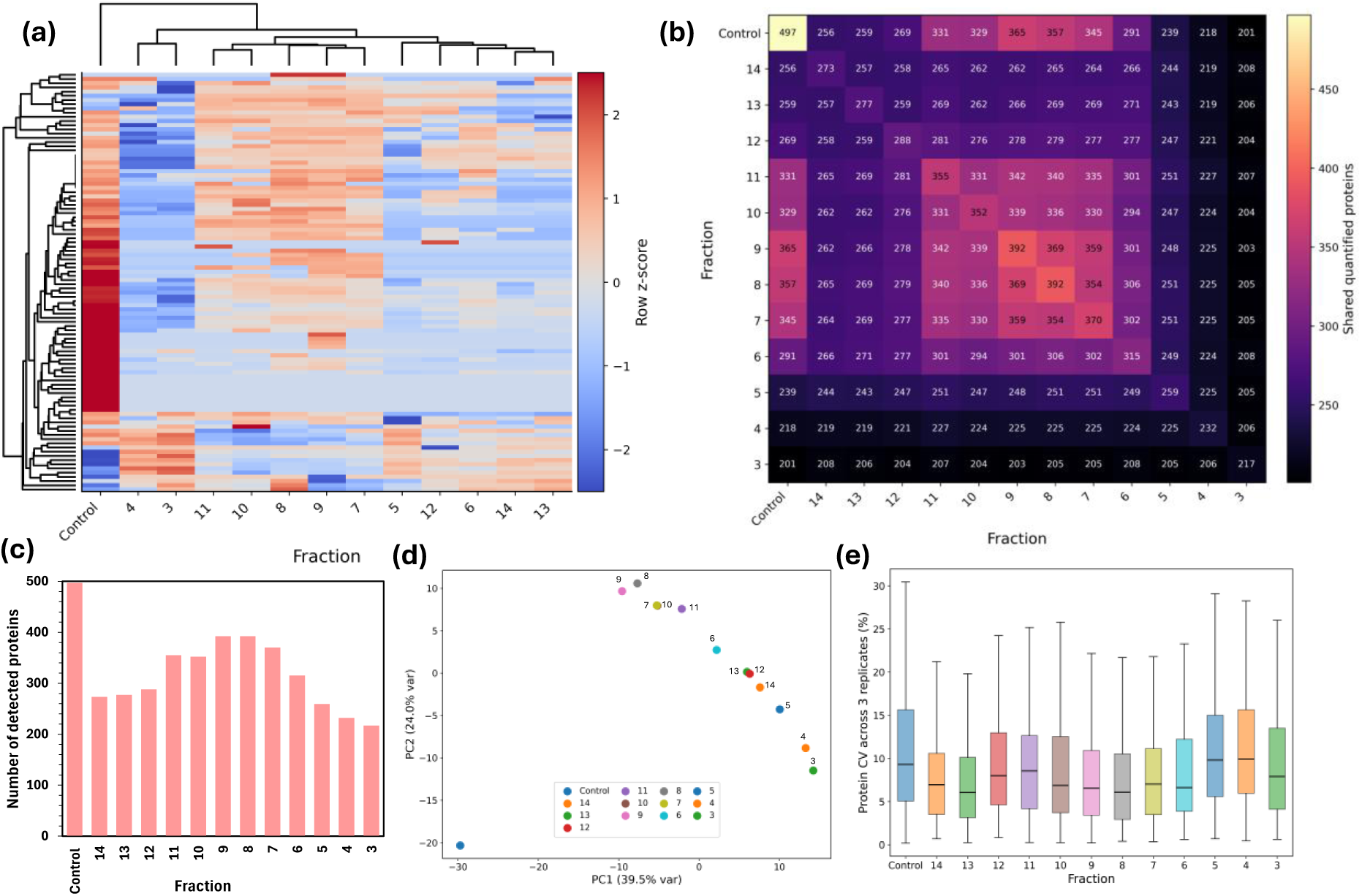
(a) Clustered heatmap of normalized protein abundances across control and fractions 14 to 3, (b) pairwise shared-protein count heatmap across fractions, (c) number of detected proteins per fraction, (d) PCA of fraction-level proteomic profiles, and (e) replicate reproducibility shown as protein-level coefficient of variation across fractions.

This variation is also evident in the clustered abundance heatmap, which shows differences in protein abundance patterns across fractions, and in the shared-protein overlap matrix, which shows how protein sharing changes between fractions (**Figures 7a** and **7b**). The clustered heatmap shows that proteins segregate into coordinated abundance patterns across the fraction series, with fractions arranging in an ordered manner rather than forming a few repeated or duplicated blocks. Moreover, the shared-protein heatmap (**Figure 7b**) demonstrates that the highest overlap occurs between nearby fractions, whereas overlap decreases progressively as the physical distance between fractions increases. For example, neighboring fractions in the middle portion of the column share substantially more proteins than distant upper-versus-lower comparisons, indicating that the proteome is remodeled gradually along the MagLev column. The unfractionated control remains the most distinct sample overall, while the internal fraction series form a structured proteomic trend. PCA further supports these results by showing that the fractions follow an ordered trend along the MagLev column rather than forming a few clearly separated clusters (**Figure 7d**). The control separates clearly from the fractionated samples, indicating that MagLev fractionation substantially restructures the proteome relative to the initial PC batch. Within the fraction series, the PCA shows gradual changes from the upper to the lower fractions, consistent with the abundance heatmap and overlap analysis, and supports a smooth but structured fraction-dependent transition.

The biological meaning of this redistribution becomes clearer when comparing the proteins with the strongest trajectory changes across fractions 14 to 3. The most dynamic proteins included KRT9, PGK1, an Ig-like domain-containing protein, CCT3, CORO1C, TUBB1, ADH4, TLN1, KRT14, KRT1, PROS1, and STOM (**Figure 8a**). These proteins point to a mixture of structural/keratin-associated, cytoskeletal, protein-folding, metabolic/enzymatic, and plasma- or extracellular-associated functions.^60^ In particular, the prominence of KRT9, KRT14, KRT1, STOM, TLN1, and TUBB1 indicates that structural and membrane-associated protein classes contribute strongly to the fraction-dependent differences, while CCT3 and CORO1C indicate remodeling of cytoskeletal organization and intracellular protein-handling processes (**Figure 8b**). At the same time, PGK1 and ADH4 demonstrate the contribution of metabolic enzymes, whereas PROS1 indicates that extracellular or plasma-associated signals remain represented among the changing proteins. Importantly, these proteins do not simply show monotonic high-to-low or low-to-high shifts; instead, many exhibit fraction-specific peaks and troughs, demonstrating the localized enrichment of distinct biochemical states in different regions of the high-sensitivity MagLev column. These results indicate that the apparent PC is not compositionally uniform even among nominally identical NPs, but instead consists of distinct subpopulations that are averaged out in bulk measurements and therefore needs to be carefully considered in nanomedicines.

**Figure 8:**
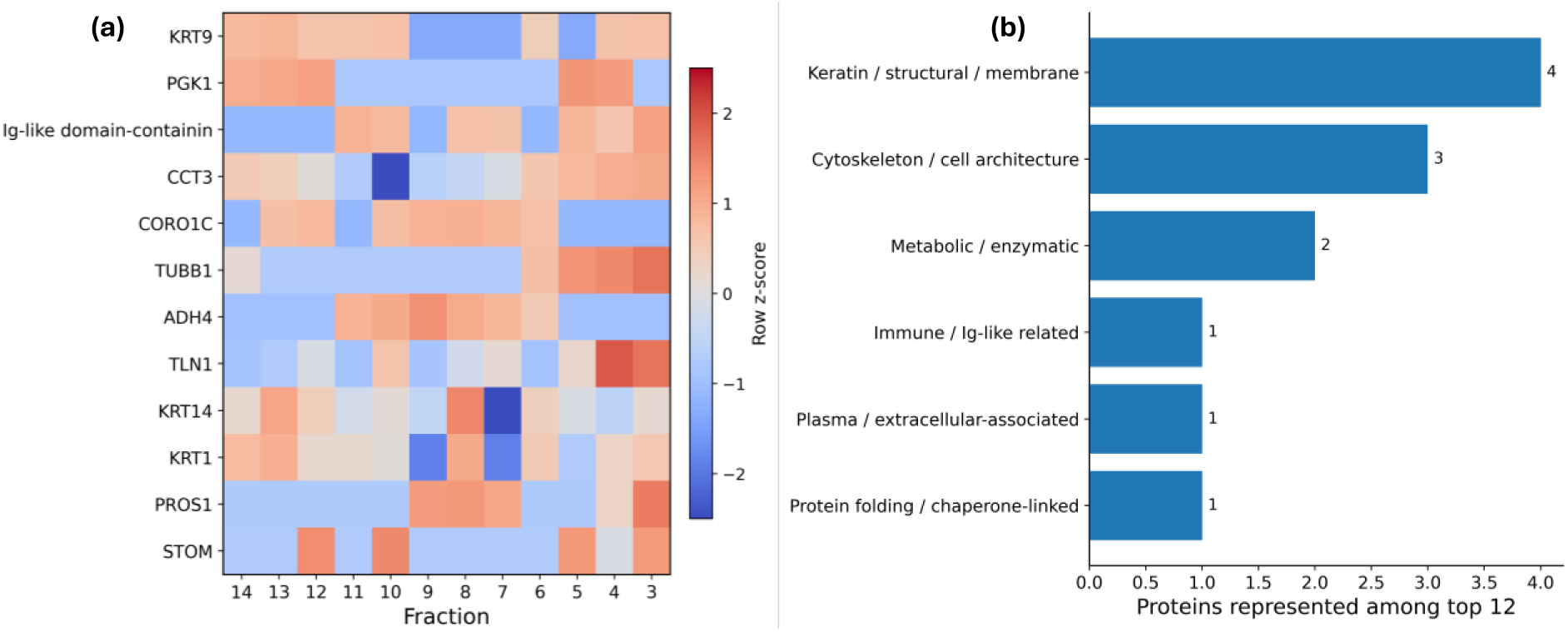
(a) Heatmap of the top 12 proteins with the strongest fraction-dependent abundance trajectories across fractions 14 to 3 and (b) functional theme summary showing the major biological classes represented among the top-changing proteins.

The proteomics data indicates that high-sensitivity MagLev does not merely separate samples by differences in total protein abundance, but instead reflects a continuous and biologically meaningful redistribution of protein identity across fractions. In other words, there is a top-to-bottom gradient of changing protein composition in which adjacent fractions remain related while gradually diverging over larger distances. This pattern is consistent with the ability of the high-sensitivity MagLev system to detect subtle differences in PC composition due to increased resolution compared with standard Maglev and to translate those differences into measurable proteomic heterogeneity. ^3^ In this context, the fraction series can be understood as a high-resolution map of related but distinct proteomic states, with the middle fractions showing the broadest protein coverage and the strongest overall overlap, and the upper and lower fractions capturing more specialized subsets of the proteome.

Beyond the number of proteins and boundary statistics, the gene ontology (GO) analysis revealed distinct differences in PC composition across the various fractions, providing a biologically meaningful interpretation of the fractionation process (**Table S1**). Across the MagLev fraction series, the proteomic composition changes not only in abundance and overlap, but also in biological identity, revealing a meaningful redistribution of protein classes from the upper to the lower regions of the column. The upper fractions are relatively richer in proteins associated with extracellular and plasma-related functions, including immune- and adhesion-linked signatures, consistent with retention of more circulating or surface-accessible protein populations near the top of the MagLev path. As the sample moves downward, the proteome becomes increasingly enriched in proteins linked to structural organization, membrane association, cytoskeletal remodeling, and intracellular biochemical processes, indicating a shift toward more cell-architecture- and protein-organization-related signatures. This trend is supported by the prominence of keratin- and membrane-associated proteins such as KRT9, KRT14, KRT1, and STOM, together with proteins involved in cytoskeletal structure and organization such as TLN1, TUBB1, and CORO1C. In parallel, the appearance of PGK1 and ADH4 indicates that metabolic enzymes are also redistributed across the fraction series, while proteins such as CCT3 suggest contributions from protein-folding and proteostasis-related pathways (**Figure 8**). The presence of PROS1 and an Ig-like domain-containing protein further shows that extracellular/plasma-associated and immune-related components remain represented among the proteins that change most strongly across fractions.

Taken together, these patterns indicate that movement from the upper to the lower MagLev fractions is associated with a transition from a more extracellular/plasma/immune-associated proteomic signature toward a more structural/membrane/cytoskeletal/metabolic signature. Importantly, this change is not a simple binary switch and does not reflect complete replacement of one protein population by another. In fact, the data suggest a trend in which the MagLev column resolves a continuous but biologically meaningful redistribution of related protein populations, with neighboring fractions remaining proteomically similar while gradually diverging across the full top-to-bottom range. In pathway-level terms, the dominant biological themes implicated by the changing proteins include cell architecture and cytoskeletal organization, membrane-associated structure, protein folding/proteostasis, metabolic enzyme activity, and extracellular/immune-associated biology. Therefore, we may conclude that the high-sensitivity MagLev platform separates the sample not only by differences in protein amount, but by differences in protein type, functional role, and associated biological state.

To complement the global fraction-level analyses, the strongest-changing proteins were studied separately to identify the biological classes that contribute most prominently to the fraction-dependent redistribution. The top-changing set (**Figure 8**) was dominated by keratin/structural/membrane-associated, cytoskeletal/cell-architecture, metabolic/enzymatic, protein-folding, and smaller extracellular/immune-related signatures. In particular, KRT9, KRT14, KRT1, and STOM highlighted strong structural and membrane-associated contributions, whereas TLN1, TUBB1, and CORO1C pointed to cytoskeletal organization. Additional contributions from PGK1, ADH4, CCT3, PROS1, and an Ig-like domain-containing protein indicated representation of metabolic, proteostasis-related, extracellular, and immune-linked functions. The results suggest that the fraction series differs not only in protein abundance, but also in the biological classes of proteins that become enriched across the column.

These findings are important because they show that even when NPs are generated under the same experimental conditions and would typically be regarded as carrying the same corona, the PC composition is in fact heterogeneous at the subpopulation level. Bulk proteomics tends to collapse this complexity into a single averaged composition, potentially masking meaningful differences in protein identity, functional class, and associated biological state.^59^ In contrast, high-sensitivity MagLev reveals that the PC is distributed across fractions with distinct proteomic signatures, indicating that even identical NPs can form different PC compositions.^59^ This has direct implications for NPs characterization and nanomedicine, since biological identity, transport, targeting, and downstream cellular interactions may depend not only on the average PC composition, but also on the heterogeneity of PC-bearing subpopulations within the sample.

## Conclusions

In summary, we developed a high-sensitivity MagLev platform that enables fractionation of nominally identical PC-coated NPs with density resolution substantially beyond that of standard MagLev systems. This approach revealed that apparently uniform NPs populations are in fact compositionally heterogeneous, with distinct PC-bearing subpopulations distributed across the MagLev column. Fraction-resolved proteomic analysis showed that the PC composition does not change only in total protein abundance, but also in protein identity, overlap, and biological function, producing a structured top-to-bottom redistribution of proteomic signatures across fractions. Our findings show progressive changes in structural, cytoskeletal, metabolic, extracellular, and immune-associated protein classes, demonstrating that pooled or bulk proteomics can mask important subpopulation-level differences in PC composition. Overall, these findings establish high-sensitivity MagLev as a simple, label-free analytical framework for resolving hidden PC heterogeneity and for studying NPs-biomolecule interactions with a level of compositional resolution that is not accessible through conventional bulk analysis. This capability may be broadly useful for improving NPs characterization, understanding nano-bio interactions, and advancing the rational design and clinical translation of nanomedicines.

## Associated content

### Supporting Information

Supporting figures/tables and LC-MS/MS analyses. This material is available free of charge via the internet at http://pubs.acs.org.

## Notes

The authors declare no competing financial interest.

## Acknowledgments

All authors acknowledge financial support from National Institutes of Health (grant number R03EB034817).

---

**Figure S1:**
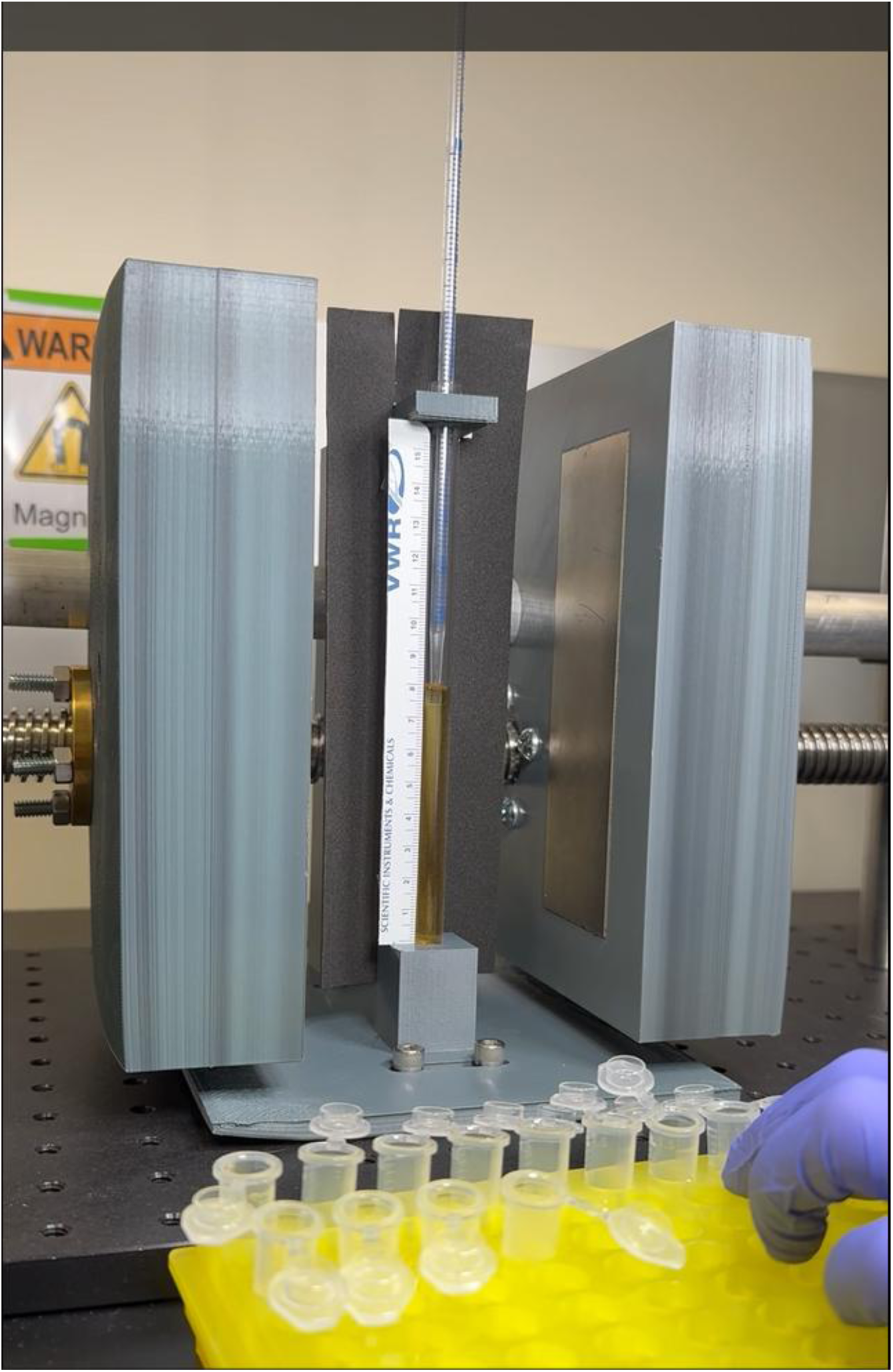
Photograph showing the extraction of different fractions from top to bottom of the high-sensitivity MagLev system using a micropipette attached to a 10 ml glass Pasteur pipette, with fractions collected at 1 cm intervals along the MagLev column and transferred to low binding Eppendorf tubes.

**Figure S2:**
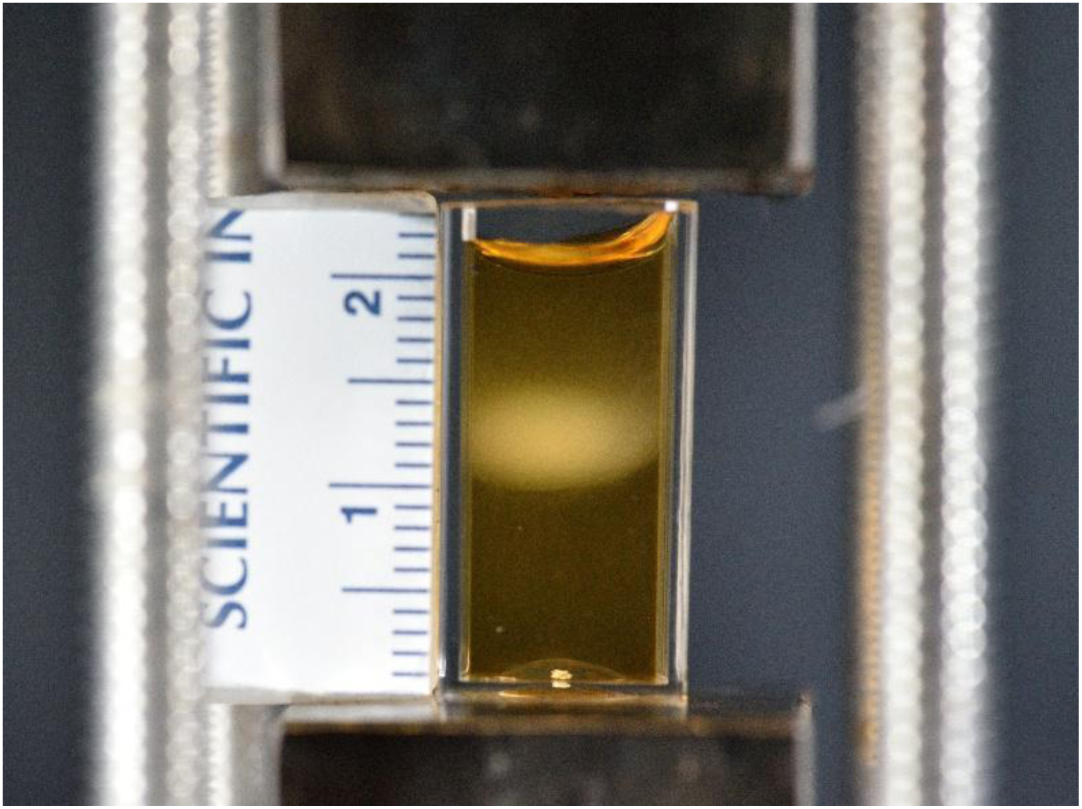
Levitation image of the PC-coated NPs in 0.125 mg/ml SPIONs in a standard MagLev system.

**Table S1:**
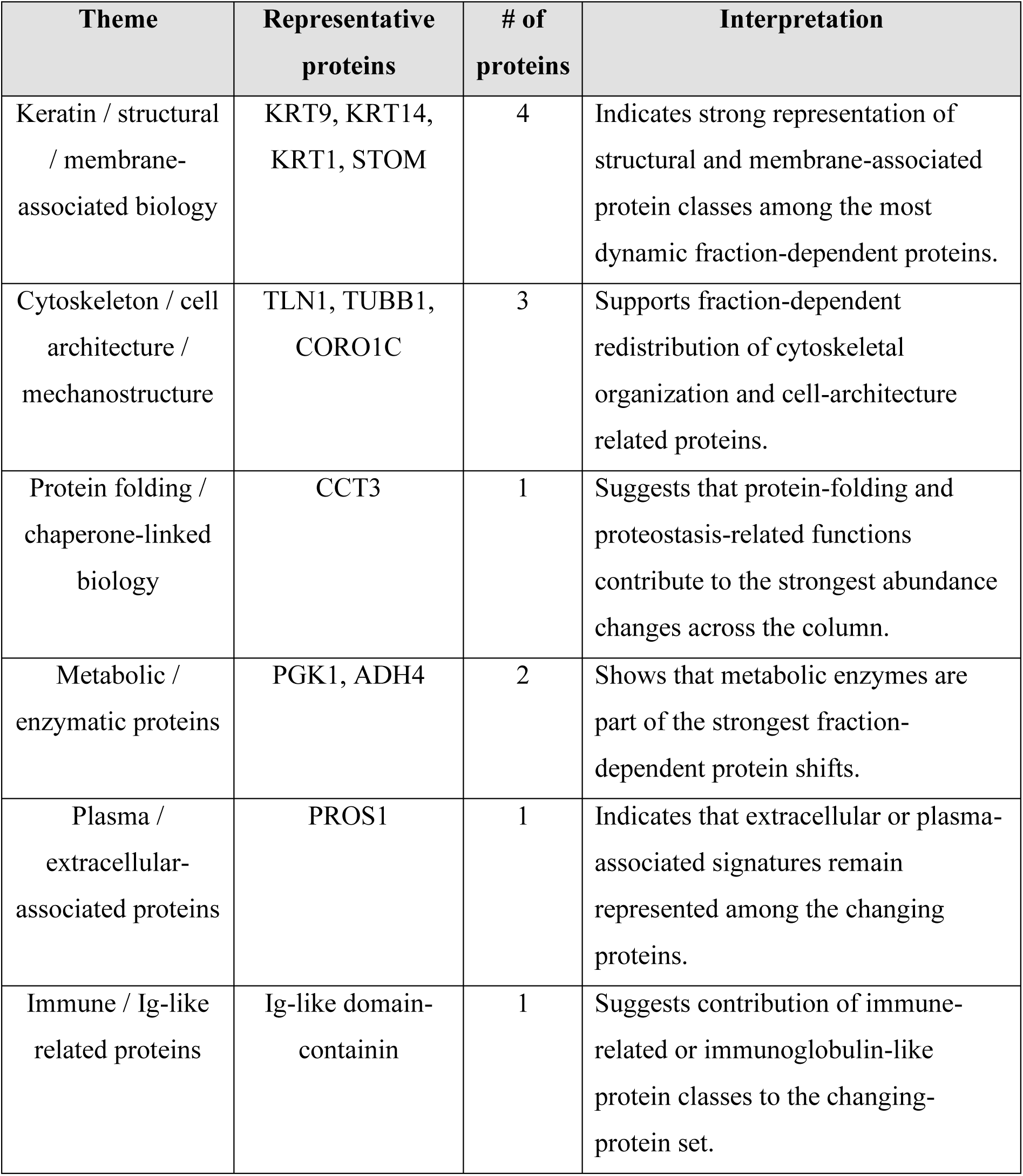
Representative GO/pathway-related protein themes identified across the changing-protein dataset, with example proteins and corresponding biological interpretation. Analysis of the most dynamic proteins across the high-sensitivity MagLev fractions indicates prominent representation of structural, cytoskeletal, membrane-associated, metabolic, and selected extracellular protein classes, rather than a purely plasma- or immune-dominated signature alone. In particular, keratin- and membrane-associated proteins such as KRT9, KRT14, KRT1, and STOM, together with cytoskeletal and cell-architecture proteins such as TLN1, TUBB1, and CORO1C, represent the largest component of the top-changing set. This suggests that the strongest fraction-dependent differences are related prominently to structural organization and protein assemblies associated with membrane or cytoskeletal states. Moreover, metabolic enzymes such as PGK1 and ADH4, the chaperone-linked protein CCT3, the plasma-associated protein PROS1, and an Ig-like domain-containing protein indicate that the fractionation also captures contributions from metabolism, proteostasis, extracellular biology, and immune-related signatures. The findings reveal that the high-sensitivity MagLev fractions differ not only in protein number and overlap, but also in protein class, with the top-changing proteins emphasizing structural and cytoskeletal remodeling alongside metabolic and extracellular-associated components.

## References

1. Monopoli, M. P.; Åberg, C.; Salvati, A.; Dawson, K. A., Biomolecular coronas provide the biological identity of nanosized materials. Nature nanotechnology 2012, 7 (12), 779–786.

2. Tenzer, S.; Docter, D.; Kuharev, J.; Musyanovych, A.; Fetz, V.; Hecht, R.; Schlenk, F.; Fischer, D.; Kiouptsi, K.; Reinhardt, C.; Landfester, K.; Schild, H.; Maskos, M.; Knauer, S. K.; Stauber, R. H., Rapid formation of plasma protein corona critically affects nanoparticle pathophysiology. Nature Nanotechnology 2013, 8 (10), 772–781.

3. Ashkarran, A. A.; Gharibi, H.; Voke, E.; Landry, M. P.; Saei, A. A.; Mahmoudi, M., Measurements of heterogeneity in proteomics analysis of the nanoparticle protein corona across core facilities. Nature Communications 2022, 13 (1), 6610.

4. Sheibani, S.; Basu, K.; Farnudi, A.; Ashkarran, A.; Ichikawa, M.; Presley, J.; Bui, K.; Ejtehadi, M. R.; Vali, H.; Mahmoudi, M., Nanoscale Characterization of the Biomolecular Corona by Cryo-Electron Microscopy, Cryo-Electron Tomography, and Image Simulation. Nature Communications 2021, 12, 573.

5. Mahmoudi, M.; Landry, M. P.; Moore, A.; Coreas, R., The protein corona from nanomedicine to environmental science. Nature Reviews Materials 2023.

6. Wang, J.; Jensen, U. B.; Jensen, G. V.; Shipovskov, S.; Balakrishnan, V. S.; Otzen, D.; Pedersen, J. S.; Besenbacher, F.; Sutherland, D. S., Soft interactions at nanoparticles alter protein function and conformation in a size dependent manner. Nano Letters 2011, 11 (11), 4985–4991.

7. Kelly, P. M.; Åberg, C.; Polo, E.; O’Connell, A.; Cookman, J.; Fallon, J.; Krpetić, Ž.; Dawson, K. A., Mapping protein binding sites on the biomolecular corona of nanoparticles. Nature Nanotechnology 2015, 10 (5), 472–479.

8. Ashkarran, A. A.; Sharifi, S.; LaRock, C. N.; Mahmoudi, M., Improved Methodology for Evaluating Nanomedicine Antibacterial Properties in Biological Fluids. ACS Pharmacology & Translational Science 2024, 7 (3), 855–862.

9. Hajipour, M. J.; Safavi-Sohi, R.; Sharifi, S.; Mahmoud, N.; Ashkarran, A. A.; Voke, E.; Serpooshan, V.; Ramezankhani, M.; Milani, A. S.; Landry, M. P.; Mahmoudi, M., An Overview of Nanoparticle Protein Corona Literature. Small 2023, *n/a* (n/a), 2301838.

10. Zhang, Y.; Zhang, L.; Cai, C.; Zhang, J.; Lu, P.; Shi, N.; Zhu, W.; He, N.; Pan, X.; Wang, T.; Feng, Z., In situ study of structural changes: Exploring the mechanism of protein corona transition from soft to hard. Journal of Colloid and Interface Science 2024, 654, 935–944.

11. Blume, J. E.; Manning, W. C.; Troiano, G.; Hornburg, D.; Figa, M.; Hesterberg, L.; Platt, T. L.; Zhao, X.; Cuaresma, R. A.; Everley, P. A., Rapid, deep and precise profiling of the plasma proteome with multi-nanoparticle protein corona. Nature communications 2020, 11 (1), 3662.

12. Ashkarran, A. A.; Gharibi, H.; Grunberger, J. W.; Saei, A. A.; Khurana, N.; Mohammadpour, R.; Ghandehari, H.; Mahmoudi, M., Sex-Specific Silica Nanoparticle Protein Corona Compositions Exposed to Male and Female BALB/c Mice Plasmas. ACS Bio & Med Chem Au 2023, 3 (1), 62–73.

13. Hajipour, M. J.; Laurent, S.; Aghaie, A.; Rezaee, F.; Mahmoudi, M., Personalized protein coronas: A “key” factor at the nanobiointerface. Biomaterials Science 2014, 2 (9), 1210–1221.

14. Settanni, G.; Zhou, J.; Suo, T.; Schöttler, S.; Landfester, K.; Schmid, F.; Mailänder, V., Protein corona composition of poly(ethylene glycol)-and poly (phosphoester)-coated nanoparticles correlates strongly with the amino acid composition of the protein surface. Nanoscale 2017, 9 (6), 2138–2144.

15. Saorin, A.; Martinez-Serra, A.; Henry, M.; Meleady, P.; Monopoli, M. P., Mass Spectrometry Proteomics of the Nanoparticle Corona Is Highly Dependent on Sample Preparation Protocol. PROTEOMICS 2026, *n/a* (n/a), e70118.

16. Ashkarran, A. A., Decentralized nanoparticle protein corona analysis may misconduct biomarker discovery. Nano Today 2025, 62, 102745.

17. Mahmoudi, M., Debugging Nano–Bio Interfaces: Systematic Strategies to Accelerate Clinical Translation of Nanotechnologies. Trends in Biotechnology 2018, 36 (8), 755–769.

18. Mahmoudi, M.; Bertrand, N.; Zope, H.; Farokhzad, O. C., Emerging understanding of the protein corona at the nano-bio interfaces. Nano Today 2016, 11 (6), 817–832.

19. Mahmoudi, M.; Saeedi-Eslami, S. N.; Shokrgozar, M. A.; Azadmanesh, K.; Hassanlou, M.; Kalhor, H. R.; Burtea, C.; Rothen-Rutishauser, B.; Laurent, S.; Sheibani, S.; Vali, H., Cell “vision”: Complementary factor of protein corona in nanotoxicology. Nanoscale 2012, 4 (17), 5461–5468.

20. Zhang, P.; Cao, M.; Chetwynd, A. J.; Faserl, K.; Abdolahpur Monikh, F.; Zhang, W.; Ramautar, R.; Ellis, L.-J. A.; Davoudi, H. H.; Reilly, K.; Cai, R.; Wheeler, K. E.; Martinez, D. S. T.; Guo, Z.; Chen, C.; Lynch, I., Analysis of nanomaterial biocoronas in biological and environmental surroundings. Nature Protocols 2024, 19 (10), 3000–3047.

21. Dabbagh, S. R.; Alseed, M. M.; Saadat, M.; Sitti, M.; Tasoglu, S., Biomedical Applications of Magnetic Levitation. Advanced NanoBiomed Research 2022, 2 (3), 2100103.

22. Alseed, M. M.; Rahmani Dabbagh, S.; Zhao, P.; Ozcan, O.; Tasoglu, S., Portable magnetic levitation technologies. 2021, 10 (2), 109–121.

23. Amin, R.; Knowlton, S.; Yenilmez, B.; Hart, A.; Joshi, A.; Tasoglu, S., Smart-phone attachable, flow-assisted magnetic focusing device. RSC Advances 2016, 6 (96), 93922–93931.

24. Tasoglu, S.; Yu, C. H.; Liaudanskaya, V.; Guven, S.; Migliaresi, C.; Demirci, U., Magnetic Levitational Assembly for Living Material Fabrication. Advanced healthcare materials 2015, 4 (10), 1469–76, 1422.

25. Anil-Inevi, M.; Sarigil, O.; Unal, Y. C.; Tekin, H. C.; Mese, G.; Ozcivici, E., Magnetic levitation-based determination of single-nuclei density. Biomaterials Advances 2026, 180, 214581.

26. Sarigil, O.; Anil Inevi, M.; Inal, E.; Tosunoglu, P. B.; Yalcin Ozuysal, O.; Tekin, H. C.; Mese, G.; Ozcivici, E., Magnetic levitation-assisted biofabrication and 3D culture of adipocytes, endothelial cells, and macrophages. Sensors and Actuators B: Chemical 2026, 457, 139724.

27. Karakuzu, B.; Anil İnevi, M.; Tarim, E. A.; Sarigil, O.; Guzelgulgen, M.; Kecili, S.; Cesmeli, S.; Koc, S.; Baslar, M. S.; Oksel Karakus, C.; Ozcivici, E.; Tekin, H. C., Magnetic levitation-based miniaturized technologies for advanced diagnostics. Emergent Materials 2024, 7 (6), 2323–2348.

28. Delikoyun, K.; Yaman, S.; Yilmaz, E.; Sarigil, O.; Anil-Inevi, M.; Telli, K.; Yalcin-Ozuysal, O.; Ozcivici, E.; Tekin, H. C., HologLev: A Hybrid Magnetic Levitation Platform Integrated with Lensless Holographic Microscopy for Density-Based Cell Analysis. ACS Sensors 2021, 6 (6), 2191–2201.

29. Ramarao, M.; Garcia-Gradilla, V.; Burcin Unlu, M.; Davis, R. W.; Gozde Durmus, N., Dynamic and precise electromagnetic levitation of single cells. Proceedings of the National Academy of Sciences 2025, 122 (37), e2512246122.

30. Yaman, S.; Devoe, T.; Aygun, U.; Parlatan, U.; Bobbili, M. R.; Karim, A. H.; Grillari, J.; Durmus, N. G., EV-Lev: extracellular vesicle isolation from human plasma using microfluidic magnetic levitation device. Lab on a Chip 2025, 25 (6), 1439–1451.

31. Abrahamsson, C. K.; Nagarkar, A.; Fink, M. J.; Preston, D. J.; Ge, S.; Bozenko, J. S.; Whitesides, G. M., Analysis of Powders Containing Illicit Drugs Using Magnetic Levitation. Angewandte Chemie - International Edition 2020, 59 (2), 874–881.

32. Durmus, N. G.; Tekin, H. C.; Guven, S.; Sridhar, K.; Yildiz, A. A.; Calibasi, G.; Ghiran, I.; Davis, R. W.; Steinmetz, L. M.; Demirci, U., Magnetic levitation of single cells. Proceedings of the National Academy of Sciences of the United States of America 2015, 112 (28), E3661–E3668.

33. Parfenov, V. A.; Khesuani, Y. D.; Petrov, S. V.; Karalkin, P. A.; Koudan, E. V.; Nezhurina, E. K.; Pereira, F. D. A. S.; Krokhmal, A. A.; Gryadunova, A. A.; Bulanova, E. A.; Vakhrushev, I. V.; Babichenko, I. I.; Kasyanov, V.; Petrov, O. F.; Vasiliev, M. M.; Brakke, K.; Belousov, S. I.; Grigoriev, T. E.; Osidak, E. O.; Rossiyskaya, E. I.; Buravkova, L. B.; Kononenko, O. D.; Demirci, U.; Mironov, V. A., Magnetic levitational bioassembly of 3D tissue construct in space. Science Advances 2020, 6 (29).

34. Tocchio, A.; Durmus, N. G.; Sridhar, K.; Mani, V.; Coskun, B.; El Assal, R.; Demirci, U., Magnetically Guided Self-Assembly and Coding of 3D Living Architectures. Advanced Materials 2018, 30 (4).

35. Anil-Inevi, M.; Delikoyun, K.; Mese, G.; Tekin, H. C.; Ozcivici, E., Magnetic levitation assisted biofabrication, culture, and manipulation of 3D cellular structures using a ring magnet based setup. Biotechnology and Bioengineering 2021, 118 (12), 4771–4785.

36. Sarigil, O.; Anil-Inevi, M.; Firatligil-Yildirir, B.; Unal, Y. C.; Yalcin-Ozuysal, O.; Mese, G.; Tekin, H. C.; Ozcivici, E., Scaffold-free biofabrication of adipocyte structures with magnetic levitation. Biotechnology and Bioengineering 2021, 118 (3), 1127–1140.

37. Anil-Inevi, M.; Yaman, S.; Yildiz, A. A.; Mese, G.; Yalcin-Ozuysal, O.; Tekin, H. C.; Ozcivici, E., Biofabrication of in situ Self Assembled 3D Cell Cultures in a Weightlessness Environment Generated using Magnetic Levitation. Scientific Reports 2018, 8 (1).

38. Pakpour, S.; Vojnits, K.; Alousi, S.; Khalid, M. F.; Fowler, J. D.; Wang, Y.; Tan, A. M.; Lam, M. I.; Zhao, M.; Calderon, E.; Luka, G. S.; Hoorfar, M.; Kazemian, N.; Isazadeh, S.; Ashkarran, A. A.; Runstadler, J. A.; Mahmoudi, M., Magnetic Levitation System Isolates and Purifies Airborne Viruses. ACS Nano 2023, 17 (14), 13393–13407.

39. Yaman, S.; Tekin, H. C., Magnetic Susceptibility-Based Protein Detection Using Magnetic Levitation. Analytical Chemistry 2020, 92 (18), 12556–12563.

40. Sarigil, O.; Anil-Inevi, M.; Yilmaz, E.; Mese, G.; Tekin, H. C.; Ozcivici, E., Label-free density-based detection of adipocytes of bone marrow origin using magnetic levitation. The Analyst 2019, 144 (9), 2942–2953.

41. Ugur, A.; Sena, Y.; Ugur, P.; Naside Gozde, D. In Nanoscale detection of EpCAM-positive extracellular vesicles using magnetic beads and interferometric scattering microscopy, Proc.SPIE, 2025; p 135840B.

42. Tasoglu, S.; Khoory, J. A.; Tekin, H. C.; Thomas, C.; Karnoub, A. E.; Ghiran, I. C.; Demirci, U., Levitational Image Cytometry with Temporal Resolution. Advanced Materials 2015, 27 (26), 3901–3908.

43. Amin, R.; Knowlton, S.; Dupont, J.; Bergholz, J. S.; Joshi, A.; Hart, A.; Yenilmez, B.; Yu, C. H.; Wentworth, A.; Zhao, J. J.; Tasoglu, S., 3D-Printed Smartphone-Based Device for Label-Free Cell Separation. Journal of 3D Printing in Medicine 2017, 1 (3), 155–164.

44. Kecili, S.; Yilmaz, E.; Ozcelik, O. S.; Anil-Inevi, M.; Gunyuz, Z. E.; Yalcin-Ozuysal, O.; Ozcivici, E.; Tekin, H. C., μDACS platform: A hybrid microfluidic platform using magnetic levitation technique and integrating magnetic, gravitational, and drag forces for density-based rare cancer cell sorting. Biosensors and Bioelectronics: X 2023, 15, 100392.

45. Deshmukh, S. S.; Shakya, B.; Chen, A.; Durmus, N. G.; Greenhouse, B.; Egan, E. S.; Demirci, U., Multiparametric biophysical profiling of red blood cells in malaria infection. Communications Biology 2021, 4 (1), 697.

46. Kumar, A. A.; Chunda-Liyoka, C.; Hennek, J. W.; Mantina, H.; Lee, S. Y. R.; Patton, M. R.; Sambo, P.; Sinyangwe, S.; Kankasa, C.; Chintu, C.; Brugnara, C.; Stossel, T. P.; Whitesides, G. M., Evaluation of a density-based rapid diagnostic test for sickle cell disease in a clinical setting in Zambia. PLoS ONE 2014, 9 (12).

47. Song, Y.; Huang, Y. Y.; Liu, X.; Zhang, X.; Ferrari, M.; Qin, L., Point-of-care technologies for molecular diagnostics using a drop of blood. Trends in Biotechnology 2014, 32 (3), 132–139.

48. Knowlton, S. M.; Yenilmez, B.; Amin, R.; Tasoglu, S., Magnetic Levitation Coupled with Portable Imaging and Analysis for Disease Diagnostics. 1940-087X: 2017; Vol. 120, p e55012.

49. Felton, E. J.; Velasquez, A.; Lu, S.; Murphy, R. O.; ElKhal, A.; Mazor, O.; Gorelik, P.; Sharda, A.; Ghiran, I. C., Detection and quantification of subtle changes in red blood cell density using a cell phone. Lab on a Chip 2016, 16 (17), 3286–3295.

50. Moncal, K. K.; Najmi, L. A.; Gupta, R.; Ramarao, M.; Knowles, J. W.; Park, C. Y.; Durmus, N. G., Label-Free Detection of Lipid Accumulation in Cells Using Magnetic Levitation. Advanced biology 2025, 9 (7), e2200142.

51. Ashkarran, A. A.; Gharibi, H.; Zeki, D. A.; Radu, I.; Khalighinejad, F.; Keyhanian, K.; Abrahamsson, C. K.; Ionete, C.; Saei, A. A.; Mahmoudi, M., Multi-omics analysis of magnetically levitated plasma biomolecules. Biosensors and Bioelectronics 2023, 220, 114862.

52. Ashkarran, A. A.; Olfatbakhsh, T.; Ramezankhani, M.; Crist, R. C.; Berrettini, W. H.; Milani, A. S.; Pakpour, S.; Mahmoudi, M., Evolving Magnetically Levitated Plasma Proteins Detects Opioid Use Disorder as a Model Disease. Advanced Healthcare Materials 2020, 9 (5), 1901608.

53. Ashkarran, A. A.; Suslick, K. S.; Mahmoudi, M., Magnetically Levitated Plasma Proteins. Analytical Chemistry 2020, 92 (2), 1663–1668.

54. Ashkarran, A. A.; Dararatana, N.; Crespy, D.; Caracciolo, G.; Mahmoudi, M., Mapping the heterogeneity of protein corona by: Ex vivo magnetic levitation. Nanoscale 2020, 12 (4), 2374–2383.

55. Nemiroski, A.; Kumar, A. A.; Soh, S.; Harburg, D. V.; Yu, H.-D.; Whitesides, G. M., High-Sensitivity Measurement of Density by Magnetic Levitation. Analytical Chemistry 2016, 88 (5), 2666–2674.

56. Martinez-Serra, A.; Cheeseman, J.; Saorin, A.; Soliman, M. G.; Dobricic, M.; Spencer, D. I. R.; Monopoli, M. P., Multiple Exposures of Plasma to Nanoparticles: A Novel Tool to Personalize Biomolecular Coronas and Fractionate Fluids. Analytical Chemistry 2025, 97 (27), 14132–14141.

57. Saorin, A.; Subrati, A.; Martinez-Serra, A.; Alonso, B.; Henry, M.; Meleady, P.; Moya, S. E.; Monopoli, M. P., A Redefined Protocol for Protein Corona Analysis on Graphene Oxide. ACS Nanoscience Au 2025, 5 (5), 388–397.

58. Ashkarran, A. A., Rethinking analytical approaches to nanoparticles protein corona heterogeneity: a call for innovation. Nanomedicine 2025, 20 (22), 2709–2712.

59. Ashkarran, A. A.; Gharibi, H.; Modaresi, S. M.; Saei, A. A.; Mahmoudi, M., Standardizing Protein Corona Characterization in Nanomedicine: A Multicenter Study to Enhance Reproducibility and Data Homogeneity. Nano Letters 2024, 24 (32), 9874–9881.

60. UniProt: the Universal Protein Knowledgebase in 2023. Nucleic acids research 2023, 51 (D1), D523–d531.

